# A modular lentiviral system for multiplexed gene perturbation and functional analysis suggests interdependence of hormone receptors in breast cancer growth *in vivo*

**DOI:** 10.64898/2025.12.07.692832

**Authors:** Seher Alam, Daria Matvienko, Johan Boström, Maria J. Pires, Aikaterini Papadimitropoulou, Saeed Eshtad, Kumar Sanjiv, Hong Qian, Nicholas C.K. Valerie, Cathrin Brisken, Mikael Altun

**Affiliations:** Division of Clinical Physiology, Department of Laboratory Medicine, Karolinska Institutet, Karolinska University Hospital; Huddinge, SE-141 52, Sweden; Swiss Institute for Experimental Cancer Research, School of Life Sciences, Ecole Polytechnique Fédérale de Lausanne, Lausanne, Switzerland; Science for Life Laboratory, Department of Oncology-Pathology, Karolinska Institutet, Box 1031, 171 21, Stockholm, Sweden; Center for Hematology and Regenerative Medicine (HERM), Department of Medicine Huddinge, Karolinska Institute, Stockholm, Sweden; Science for Life Laboratory, Department of Laboratory Medicine, Karolinska Institutet, Box 1031, 171 21, Stockholm, Sweden; The Breast Cancer Now Toby Robins Breast Cancer Research Centre, The Institute of Cancer Research, London, UK

**Keywords:** Genetic engineering, functional genomics, breast cancer, *in vivo*, multiplexing, lentiviral vectors

## Abstract

Precise and flexible control of gene expression is essential for dissecting gene function in complex biological systems. Although recent developments in genetic engineering and CRISPR/Cas9 technology have expanded tools for gene activation, suppression and editing, their application in physiologically relevant models remains challenging, time consuming, and expensive. Here, we present a modular, doxycycline-inducible vector system that integrates gene overexpression, shRNA-mediated knockdown, and CRISPR/Cas9-mediated regulation within a single, lentivirus-compatible system. The modular design allows rapid exchange of selection markers, epitope tags, and reporters via Gateway cloning, providing broad adaptability across experimental settings. In addition to standard fluorescent and luminescent reporters, the system includes advanced sensors, such as Fucci cell cycle reporters, to enable monitoring of cellular processes. By combining fluorescence barcoding with combinatorial genetic perturbations, the platform supports multiplexed analysis of gene function and genetic interactions through phenotypic characterization by fluorescence imaging or flow cytometry. We demonstrate its utility *in vivo* with breast cancer intraductal xenografts, which suggest that ER+ breast cancer cells (MCF7) rely on androgen (AR), estrogen (ER) and progesterone receptors (PR) for *in vivo* growth. This versatile gene perturbation system provides tight temporal control, streamlined implementation, and high-content phenotyping capacity facilitating efficient *in vitro* and *in vivo* studies while reducing the use of animals in *in vivo* validation experiments. It thus expands the experimental repertoire for dynamic, multigene interrogation in complex systems.

## Introduction

The study of biological systems has long relied on genetic engineering to explore gene function, protein biology and cellular processes. From the first recombinant plasmids to the advent of CRISPR/Cas9, the continuous development of genetic tools has not only kept pace with scientific discovery but has actively reshaped the landscape of biological research (Pacesa et al, 2024). With the rapid development of new methods, versatile genetic engineering systems have become indispensable in biological research. Whether producing recombinant proteins in Escherichia coli for structural studies or inhibitor assays, overexpressing tagged proteins in human cells to investigate localization and interaction networks, or using shRNA- or CRISPR/Cas9-mediated perturbations in animal models for phenotypic profiling, precise genetic engineering techniques are essential in all experimental workflows (Maity et al, 2023; Pacesa et al, 2024). In this study we present a combination of plasmids, interchangeable reporters and tags compatible with overexpression, shRNA-mediated gene silencing, and CRISPR/Cas9-mediated perturbation that enable genetic studies across diverse experimental systems.

In developing this series of plasmids, we integrated recent advances in genetic engineering, cell cycle reporter systems, and fluorescent marker technologies to create a comprehensive and flexible platform. Of particular note is the Gateway cloning system originally developed by Invitrogen (Katzen, 2007), the Gateway-compatible entry and destination vector collection developed by the Campeau and Kaufman labs (Campeau et al, 2009), and the lentiviral U6-driven shRNA vector architecture established by Stewart et al. (Stewart et al, 2003), with the cassette layout adapted from Cellecta. Our goal was to combine these state-of-the-art components into a unified system offering maximum flexibility in reporter configuration, tagging options, selection markers, and advanced readout capabilities.

To achieve this, we incorporated fourteen different fluorescent proteins that can be used as fusion tags or as independent reporters. Since the discovery of green fluorescent protein (GFP) in 1978 (Prendergast & Mann, 1978), iterative cycles of targeted and random mutagenesis have expanded the spectral range of excitation and emission and optimized photostability, protein half-life, and brightness (Ormö et al, 1996; Shaner et al, 2004; Ai et al, 2008; Sakaue-Sawano et al, 2008; Subach et al, 2011; Goedhart et al, 2012; Bajar et al, 2016). Some fluorescent reporters, such as iRFP670, iRFP702, iRFP720 (Shcherbakova & Verkhusha, 2013) and sfCherry (Nguyen et al, 2013), were developed by random mutagenesis and subsequent selection for desired properties. The FPBase database provides extensive information on existing fluorescent proteins and their derivation (Lambert, 2019). In addition, our system integrates three luminescent reporters, Luc2 (Kim et al, 2010), AkaLuc (Iwano et al, 2018) and NanoLuc (Hall et al, 2012) to extend detection capabilities beyond fluorescence.

To illustrate the potential of our system in a biologically relevant context, we focused on hormone-sensitive breast cancer (HSBC) as a disease model. HSBC represents the major proportion of all breast cancers and is characterized by the expression of estrogen receptors (ER) and/or progesterone receptors (PR). The growth and progression of these tumors are driven by estrogens, making them responsive to endocrine therapies that block hormone receptor signaling or suppress hormone production (McGuire et al, 1977; Fannon et al, 2001; Kohler et al, 2015; Horn & Vatten, 2017). Current standard treatments include selective estrogen receptor modulators such as tamoxifen, aromatase inhibitors, and selective estrogen receptor degraders (SERDs) (Kim & Lukong, 2025). Despite the clinical success of endocrine therapy in HSBC, resistance frequently develops, leading to disease recurrence and progression (Hanker et al, 2020). Understanding the molecular mechanisms governing hormone dependence and therapy resistance is therefore critical to improve therapeutic strategies and patient outcomes (Hanker et al, 2020). Advances in genetic engineering, including CRISPR/Cas9 genome editing and RNA interference, now provide a powerful means to dissect the regulatory networks that control hormone receptor function (Maity et al, 2023).

In addition, physiologically relevant models are essential for investigating HSBC. The mouse intraductal xenograft approach (<u>m</u>ouse <u>in</u>tra<u>d</u>uctal, MIND) has emerged as a robust system for establishing ER^+^ BC *in vivo* by recreating the native mammary microenvironment allowing studies under more clinically relevant endocrine conditions (Sflomos et al, 2016; Fiche et al, 2019; Constantin et al, 2025). Analysis of hormone response of MIND-PDX models revealed patient-specific responses as an important role for PR signaling in a subset of tumors (Scabia et al, 2022), prompting us to investigate the importance of PR and AR in addition to ER function.

To address these questions and extend experimental flexibility across different model systems, we developed a modular, doxycycline-inducible lentiviral vector platform for controlled gene expression and combinatorial perturbations. Through a series of proof-of-concept experiments, we demonstrate its utility for studying cellular function, monitoring dynamic cellular responses, and tracking cell fate across diverse experimental contexts. This includes performing multiplexing fluorescent barcoding to dissect hormone receptor dependencies and signaling crosstalk *in vitro* and *in vivo* using MIND. This adaptable and modular system provides a scalable framework for functional genomics, enabling the generation of refined preclinical models and advancing mechanistic studies of endocrine signaling and resistance.

## Results

### Development and validation of a flexible, inducible plasmid system for controlled genetic manipulation

To enable precise and versatile genetic manipulation, we developed a modular, lentivirus-compatible plasmid system that supports both inducible gene overexpression and targeted gene downregulation in mammalian cells (**Fig 1A**). Each module is compatible with constitutive or doxycycline-regulated systems, enabling tight and temporal control of expression.

**Figure 1.**
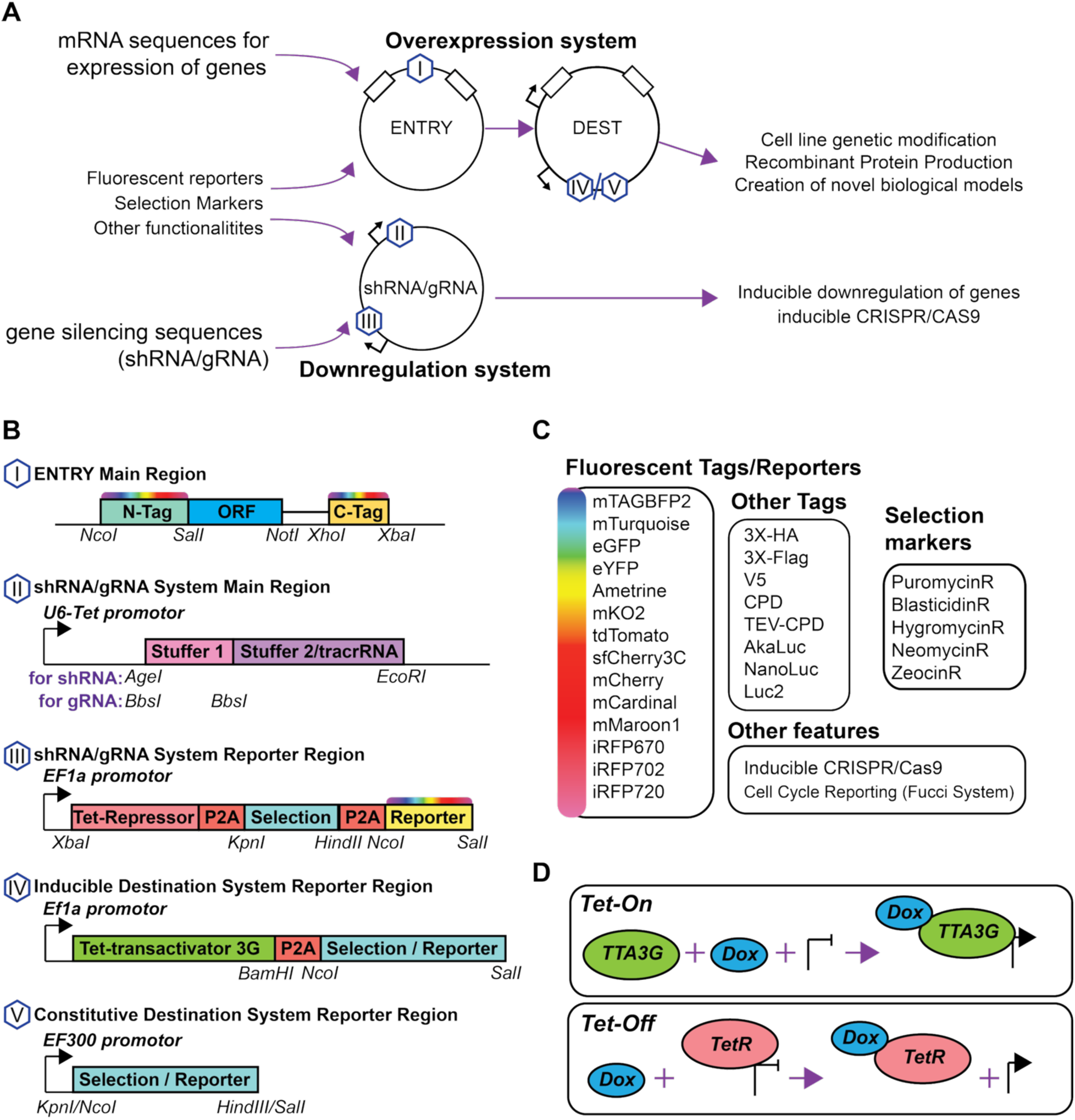
Schematic overview of the multifunctional plasmid system and its key components. **(A)** An overview of the dual-function vector system enabling gene overexpression and targeted gene knockdown in mammalian cells. **(B)** Schematic illustration of modular construct cassettes in the vector for the expression regions and the reporter. **(C)** A list of features of the plasmid system. **(D)** A schematic and comparison of the Tet-On and the Tet-Off system utilized in the plasmid system.

For gene downregulation, we designed a backbone compatible with the expression of shRNA or CRISPR/Cas9 gRNA, incorporating a general stuffer sequence flanked by cloning sites for oligo annealing (**Fig 1B**). A doxycycline-responsive U6 Tet promoter controls gRNA/shRNA transcription, which is strongly repressed by a co-expressed Tet repressor in the absence of doxycycline. It also contains an EF1α-driven tricistronic cassette containing the Tet repressor, a selectable antibiotic resistance gene or a luminescent marker, and a fluorescent or functional reporter, separated by a P2A site, a sequence that induces skipping of peptide bond formation by the ribosome to generate two separate polypeptides (Liu et al, 2017). Flanking restriction sites for each element enable the rapid exchange of selection markers and reporters.

For inducible overexpression of genes, we used a Gateway-compatible entry/destination system. The cloning strategy, based on pENTR1A/pENTR4 (Campeau et al., 2009) allows ORF insertion with customizable N- or C-terminal tags while maintaining reading frame compatibility. The target plasmids contain either constitutive (EF300) or inducible (Tet-On) promoters, and lentiviral elements enabling genomic integration for long-term expression and selection.

We integrated a library of fourteen fluorescent proteins covering the visible to near-infrared spectrum, including long Stokes shift variants (e.g. mAmetrine) to facilitate multiplex imaging (**Fig 1B and C**). In addition, we incorporated three highly sensitive luciferases (NanoLuc, AkaLuc, and Luc2) to provide complementary luminescence detection modes (**Fig 1B (I), (III)**). These reporters can be flexibly interchanged and allow one to monitor cell viability, proliferation and tumor burden in real time and in a substrate-dependent manner, while also enabling multiplexed reporting.

To enable biochemical characterization and experimentation, we integrated common non-fluorescent epitope tags such as 3×HA, 3×Flag, and V5, as well as a cysteine protease domain (CPD) tag with an optional TEV cleavage site to support solubilization, purification, and detection strategies (**Fig 1C**).

### Dual inducible CRISPR/Cas9 with synchronous activation

To achieve inducible genome editing, we co-expressed inducible Cas9 (under a Tet-On promoter) and inducible gRNA (under Tet-Off control) to form an inducible dual-vector system (**Fig 2A**). This configuration was chosen to reduce the levels of both components until needed and mitigate possible premature DNA cleavage. Because unwanted editing requires both Cas9 and gRNA to be simultaneously present above a functional threshold, and each is independently repressed through a distinct tetracycline-responsive mechanism (Tet-On for Cas9, Tet-Off for gRNA), the probability of coincident leaky expression of both components is substantially lower than for either component alone. This design logic, together with the low basal editing (∼4%) measured by ONT sequencing over 11 days without doxycycline (**Fig 2G**), indicates that this system effectively limits CRISPR/Cas9 function.

**Figure 2.**
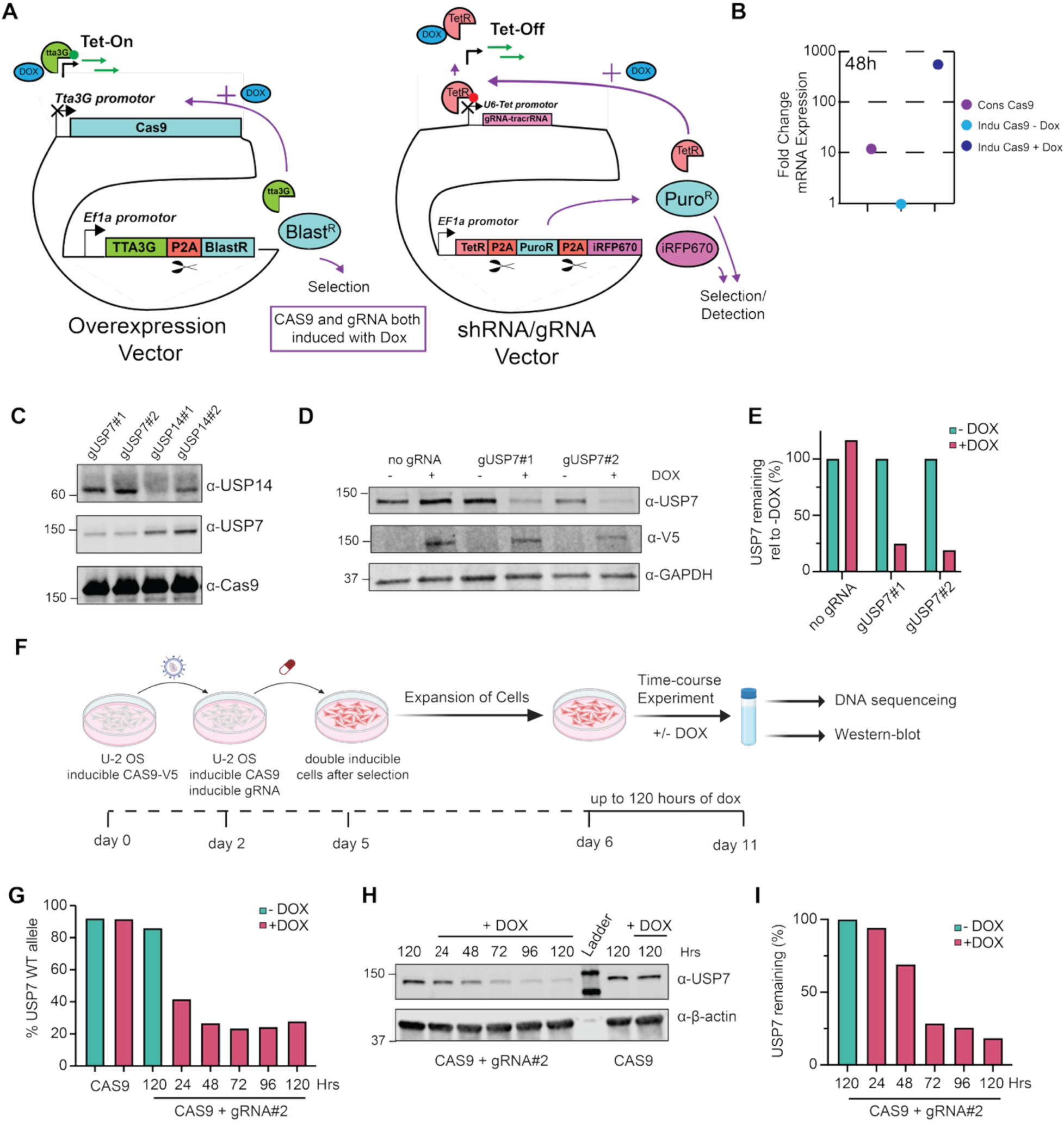
Inducible CRISPR/Cas9 system utilizing the plasmid system for tracking and functional analysis of cell populations. **(A)** Schematic overview of inducible CRISPR/CAS9 application with the plasmid system. **(B)** qPCR measurements of Cas9 mRNA comparing constitutive Cas9 to Inducible Cas9 with and without the addition of doxycycline. **(C)** Western blot of inducible CRISPR/Cas9 silencing of two deubiquitinating enzymes USP14 and USP7. **(D)** Western blot of double inducible CRISPR/Cas9 knockdown of two guides (gUSP7#1 and gUSP7#2) towards USP7 after 120 h of doxycycline induction **(E)** Quantification of Western blot shown in D. **(F)** Schematic workflow of double CRISPR/Cas9 inducible knockdown of USP7 with gRNA#2. U2OS cells expressing inducible Cas9-V5 were transduced with inducible gRNA followed by selection, and subsequent time course experiment of doxycycline induction and editing of USP7 gene up to 120 h. (**G**) Time course of USP7 editing (percentage of WT alleles) measured by ONT sequencing at indicated hours of doxycycline treatment. **(H)** Protein expression of USP7 measured by Western blot from the time course experiment described in F and G. **(I)** Quantification of USP7 protein expression from H.

Following doxycycline administration, TetR releases the U6 promoter to enable gRNA expression, while Tta3G drives robust Cas9 expression from the TRE3G promoter (**Fig 2B**). This synchronized activation of both components enables precise timing and minimizes background editing. We first validated this system using constitutive Cas9 expression and two independent inducible gRNAs targeting each of the deubiquitinating enzymes USP7 and USP14, with each gene serving as a control for knockdown of the other. Western blot analysis confirmed robust depletion of both enzymes, demonstrating the efficiency and specificity of this inducible CRISPR/Cas9 gRNA strategy (**Fig 2C**).

This inducible approach is particularly valuable when studying essential genes where leaky or constitutive editing may jeopardize cell viability. By limiting Cas9 activity with inducible control, the system enables downstream assays that could otherwise be compromised by early disruption of gene function. To evaluate the dual inducible editing, two guides towards USP7 were first assessed by Western blot 120 h after induction of inducible Cas9, showing robust depletion with both guides (**Fig 2D and E**). Furthermore, to evaluate the dynamics and leakiness of the system, ONT amplicon sequencing was performed following induction of Cas9 and one of the USP7 guides (**Fig 2F**). Editing increased rapidly after induction and reached maximal levels by 72 hours, with most edited alleles predicted to result in frameshift mutations. High editing efficiencies were maintained throughout the 120 h time course, demonstrating robust and sustained disruption of the USP7 locus **(Fig 2G).** Importantly, low background editing (4%) was observed in the non-induced sample over the 11-day period from gRNA transduction, cell expansion, and gRNA induction for 120 hours (**Fig 2G and Table S1**). This was also reflected in protein levels by Western blot (**Fig 2H and I**)

### Integration of biologically relevant reporter systems

To extend the utility of the system beyond genetic perturbations, we incorporated biologically informative reporter systems for *in vitro* and *in vivo* monitoring. Three highly sensitive luminescent reporters (NanoLuc, AkaLuc and Luc2) were integrated into the backbone at interchangeable positions, enabling real-time measurement of cell viability and cell numbers (**Fig 3A and B**). These reporters are particularly valuable for robust and quantitative endpoint readouts, longitudinal assays, as well as non-invasive xenograft studies (Sflomos et al, 2016).

**Figure 3.**
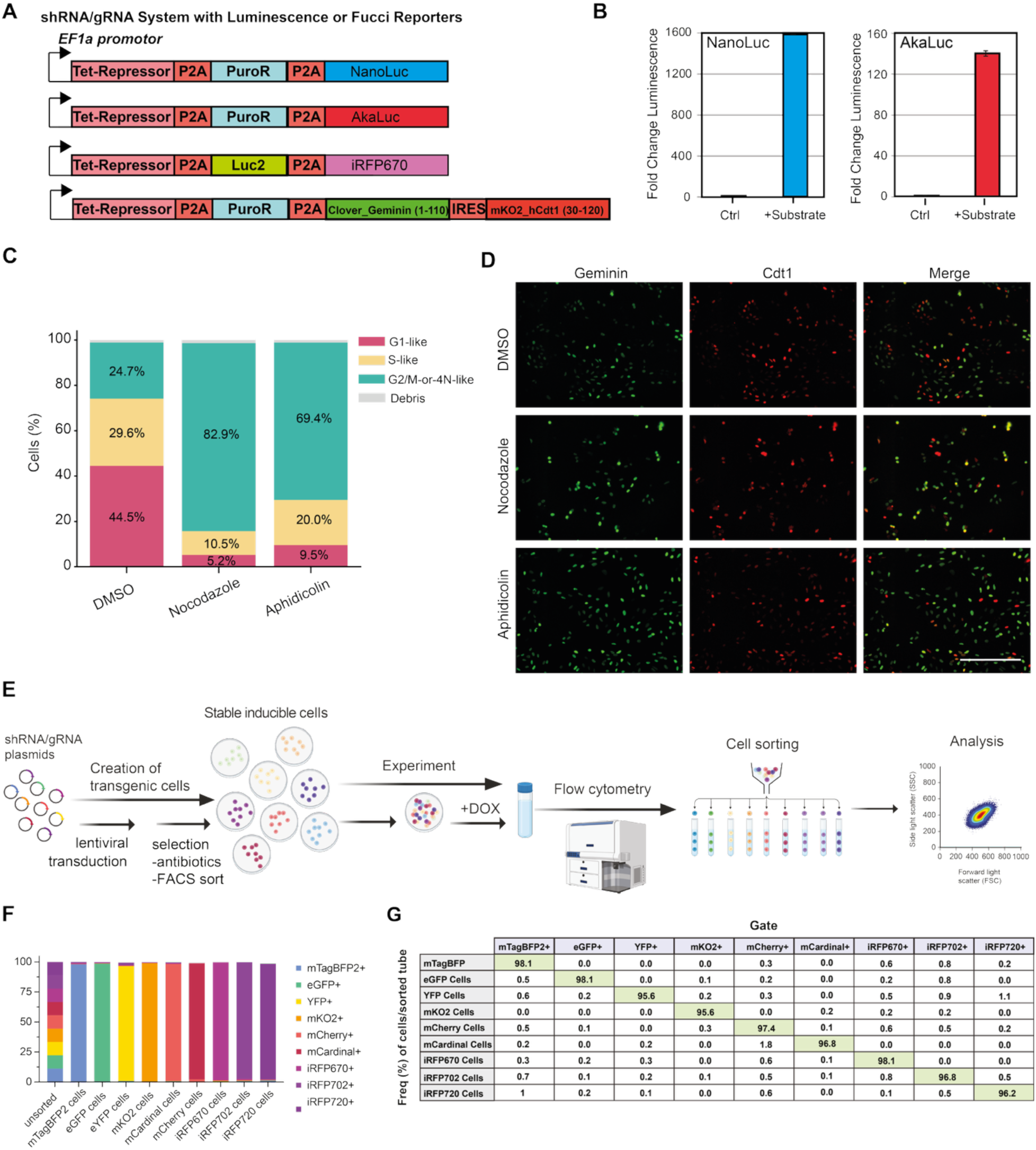
Multiplexed fluorescent barcoding and integrated luminescent and cell cycle reporters. **(A)** Schematic of lentiviral constructs with luminescent reporters (NanoLuc, AkaLuc), Luc2 with infrared fluorescent protein iRFP670 sorting and the Fucci reporter systems integrated into the plasmids. **(B)** Luminescence signal after addition of substrate relative to no substrate (Blue, NanoLuc; Red, AkaLuc). **(C)** Quantification of live cell imaging of Fucci reporters’ distribution following treatment with cell cycle arrest chemicals and DMSO control for 24 h (0.1nM Nocodazole and 1nM Aphidicolin). **(D)** Representative images of Fucci distribution in cells before and after the addition of indicated cell-cycle arresting chemicals. Scale bar = 400 µm **(E)** Schematic of fluorescence barcoding workflow. U2OS cells were transduced with nine lentiviral vectors expressing distinct fluorescent proteins, selected, pooled, treated with doxycycline, and subsequently sorted and analyzed by flow cytometry. **(F)** Bar graph showing the frequency and purity of each fluorescent population after sorting. **(G)** Table summarizing flow cytometry data used to generate the bar graph in panel F.

In the interest of general cell biology, we integrated a modified Fucci system to monitor cell cycle progression. This reporter uses fluorescently labeled fragments of the oscillating proteins hCdt1 and hGeminin to discern the G1 and S/G2/M phases, respectively (Sakaue-Sawano et al, 2008) (**Fig 3A**). After treatment with cell cycle disrupting agents (nocodazole and aphidicolin), we observed the expected fluorescence shifts, confirming the functionality of the reporters and their ability to reflect changes in cell cycle progression (**Fig 3C and D**) (Boström et al, 2017). This setup enables real-time coupling of gene disruption with cell cycle analysis and may be particularly valuable for studying proliferation-related genes in primary or slow-growing cells.

### Fluorescence barcoding enables multiplexed tracking of cell populations

Multiplexed fluorescence barcoding can enable pooling of several genetically distinct cell populations within the same well, culture, or animal, so that different genotypes are compared under identical experimental conditions rather than across separate, parallel experiments. This internally controlled design reduces confounding variability arising from well-to-well, batch-to-batch, or animal-to-animal differences and increases the statistical power obtained per experiment. It is particularly valuable *in vivo*, where it substantially reduces the number of animals required, in keeping with the 3Rs principle (Tannenbaum & Bennett, 2015), and it complements pooled multiplexed perturbation strategies used *in vitro* (Adamson et al, 2016). To support high-throughput and pooled experiments, we developed a fluorescence-based barcoding strategy using a panel of nine lentiviral vectors encoding spectrally distinct fluorescent proteins. Each vector was stably integrated into U2OS cells and selected with antibiotics to ensure uniform expression.

To discern the feasibility of a 9-plex fluorescent protein readout, we pooled the individual fluorescent bar codes for flow cytometry analysis. Pooled analysis showed that each barcode could be unambiguously resolved with minimal signal overlap, achieving 95.6–98.1% purity for the entire panel when compared with individual barcodes (**Fig 3E and G**). This enabled us to track several distinct cell populations in a single experiment and to directly compare genetic perturbations under identical experimental conditions. Such barcoding strategies are especially advantageous when using resource-intensive *in vivo* models where multiple genotypes can be tested simultaneously in the same animal, within a single tumor.

### Combinatorial hormone receptor knockdown using barcoded shRNA vectors

Next, we applied our barcoded shRNA knockdown system to investigate hormone receptor signaling in HSBC. We first transduced Luc2 into ER-positive MCF7 breast cancer cells. These were then transduced with a non-targeting control shRNA (shNT) labeled with mTagBFP2 and sorted by FACS. The sorted cells were subsequently transduced with lentiviral vectors containing shRNAs targeting ER, PR, and AR — each coupled to a unique fluorescent protein (EGFP for ER, sfCherry3C for PR, and iRFP670 for AR) — and a specific antibiotic resistance cassette. (**Fig 4A-C**).

**Figure 4.**
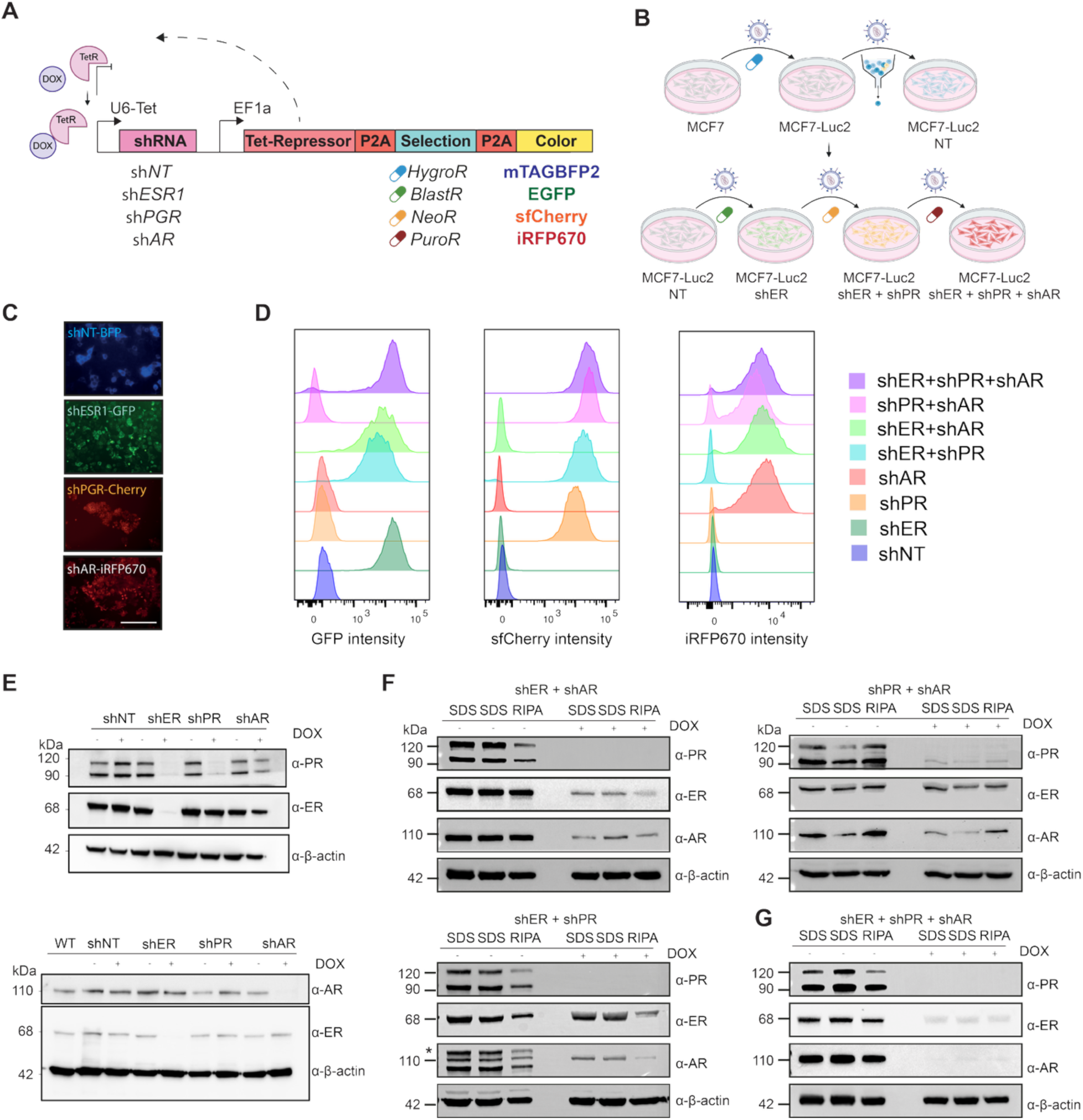
Multiplexed inducible shRNA knockdown of hormone receptors in MCF7 cells using fluorescent barcoding. **(A)** Schematic of the doxycycline-inducible multiplexed shRNA system targeting ESR1 (ER), PGR (PR), and AR. Each shRNA construct is expressed under a Tet-On promoter and coupled to a distinct fluorescent protein and antibiotic resistance marker: mTagBFP2 (NT control), GFP (ER), sfCherry3C (PR), and iRFP670 (AR). **(B)** Serial lentiviral transduction and antibiotic selection generated all single, double, and triple receptor knockdown combinations. **(C)** Microscopy image detecting fluorescent barcode after transduction and selection. Scale bar = 400µm **(D)** Histograms of fluorescence intensity showing expression of GFP, sfCherry3C, and iRFP670 in individual and combined knockdown populations. **(E)** Western blot validation of single knockdown of ER, PR, and AR protein levels following 72 hours of doxycycline treatment. β-actin was used as a loading control. **(F)** Validation of cells with double receptor knockdowns performed across replicates prepared in different lysing buffers. **(G)** Validation of triple knockdown combinations in same manner as in panel (F). * Indicates incomplete stripping of PR membrane before AR blotting.

Serial transduction was used to generate combinatorial knockdown genotypes: single (ER, PR, AR), double (ER+PR, ER+AR, PR+AR), triple (ER+PR+AR). Flow cytometry confirmed the integrity of the barcoding (**Fig 4D**), and Western blot analysis showed that the receptors were efficiently and specifically silenced by treatment with doxycycline. Additionally, silencing of ER also decreased expression of its target gene PR, as expected (Horwitz & McGuire, 1978) (**Fig 4E and G**). This approach allows for a precise analysis of combinatorial hormone-receptor interactions and, at the same time, a direct comparison between single and multi-target knockdowns.

### Multiplexed hormone receptor knockdown suppresses proliferation in vitro

To determine the functional effects of single and combined knockdowns, we examined cell proliferation after treatment with doxycycline. Viability assays showed a significant reduction in ER or PR knockdown alone (40–50% decrease), while AR depletion had minimal effects *in vitro*. Combined knockdowns further reduced viability, with triple knockdown (ER+PR+AR) showing the strongest suppression (70–80%, p < 0.001; **Fig S3A**).

Longitudinal assays over 9 days (216 hours) using cell confluence measurements confirmed these results (**Fig S3B**), and EdU incorporation assays showed that DNA synthesis was significantly impaired under all conditions with ER and/or PR depletion (**Fig S3C and D**). To assess hormone dependence, we cultured cells in hormone-depleted medium supplemented with estradiol, DHT, or promegestone. The AR-depleted cells, which did not show any proliferation deficits in full media, responded like NT cells in charcoal-stripped serum upon hormone addition, while DNA replication in ER- or PR-knockdowns remained suppressed. (**Fig S3E**). Combinations of depleted hormone receptors suppressed baseline proliferation even further while generally being non-responsive to hormone stimulation, particularly in ER+PR and ER+PR+AR knockdown cells. These results emphasize the dominant role of ER and PR in the proliferation of MCF7 cells *in vitro* and demonstrate the additive nature of multiplex knockdowns.

### Barcoded in vivo knockdowns confirm phenotypes and reduce animal use

To test the physiological relevance of the *in vitro* results *in vivo*, we intraductally injected the Luc2-labeled MCF7 cells into NSG females and validated engraftment and subsequent tumor growth via bioluminescence imaging, of mammary glands (**Fig S4A and B**). Using fluorescent barcoding, we pooled single knockdown populations for competitive *in vivo* assays. In doxycycline-treated animals, flow cytometric profiling of dissociated tumors showed selective depletion of ER-, PR-, and AR-knockdown cells (**Fig 5A-C, S5A and Table S2**).

**Figure 5.**
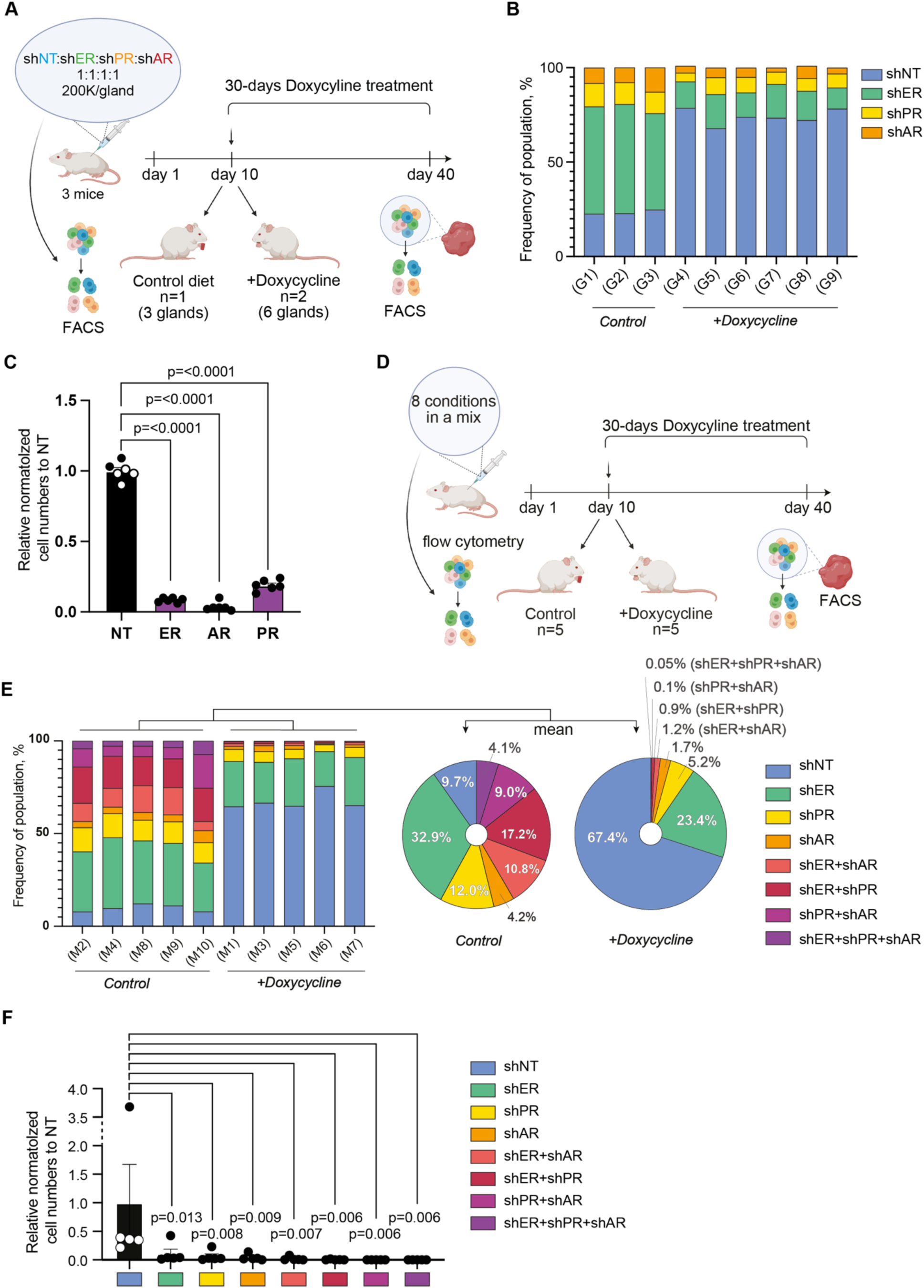
*In vivo* competition assay suggests enhanced tumor growth suppression by combined hormone receptor knockdown in MCF7 xenografts. **(A)** Schematic overview of the experimental design. MCF7 cells expressing doxycycline-inducible shRNAs targeting ER (GFP), PR (sfCherry3C), AR (iRFP670) and NT (mTagBFP2). **(B)** Number of cancer cells quantified by flow cytometry at endpoint (30 days) in doxycycline-treated (n= six glands) and the untreated control (n= three glands) mice of the different single HR knockdown cells and distribution of the different cell populations in each gland are illustrated with or without doxycycline treatment for each gland (for more details see Table S2). **(C)** Relative normalized cell numbers of the doxycycline-treated cells towards NT. Cell numbers are normalized towards the mean of each knockdown cell from the control glands, each individual datapoint are shown as circles in the graph. **(D)** Schematic overview of the experimental design. MCF7 cells expressing doxycycline-inducible shRNAs targeting ER (GFP), PR (sfCherry3C), AR (iRFP670), or combinations thereof were labeled with unique fluorescent reporters and pooled in equal ratios to create a mixed population representing all eight knockdown conditions (shNT, shER, shPR, shAR, shER+shPR, shER+shAR, shPR+shAR, shER+shPR+shAR). The cell mixture was intraductally injected into mammary glands of NSG mice. Doxycycline was administered from day 10 to day 40 post-injection to induce knockdown. At endpoint, tumors were dissociated and analyzed by flow cytometry to determine the relative abundance of each knockdown population. **(E)** Tumor composition at endpoint in doxycycline-treated (n = 5) and untreated control (n = 5) mice. Left: stacked bar plots showing the percentage of each knockdown population in individual tumors. Right: pie charts summarizing the average distribution across mice in each treatment group (for more details see Table S3). **(F)** Relative normalized cell numbers of the doxycycline-treated cells towards NT. Cell numbers are normalized towards the mean of each knockdown cell from the control animals, each individual datapoint are shown as circles in the graph. The *in vivo* statistics (**C** and **F**), the doxycycline treated animal/samples was normalized towards the corresponding average of the doxycycline untreated animal/sample for cell numbers and ANOVA with Fisher’s LSD post-hoc test towards NT. Exact p-values are indicated in the graph.

To separate the doxycycline-induced knockdown effect from potential differences in baseline engraftment or growth among the four barcoded populations, we normalized each doxycycline-treated sample to the mean cell number of the corresponding population in untreated (−Dox) control mouse glands, then expressed this value relative to the equivalently normalized non-targeting (NT) population within the same tumor (**Fig 5C and Table S2**). This approach assumes that, in the absence of knockdown, each population would engraft and grow at a rate reflected by its own untreated controls, so that any residual difference after normalization to NT reflects the knockdown effect itself rather than construct- or clone-specific variation in growth potential. Because the −Dox values are derived from separate untreated animals rather than from the same animals prior to treatment, this normalization approximates, rather than directly measures, each population’s individual growth trajectory. By this measure, ER, AR, and PR knockdown populations were reduced to 8.6% ± 0.8%, 13.4% ± 2.2%, and 19.2% ± 2.0% (mean ± SEM) of the NT population, respectively (approximately 11.6-, 7.5-, and 5.2-fold), consistent with the depletion observed by direct FACS quantification.

Finally, we injected pools containing all eight barcoded genotypes into individual mice. Initial FACS ascertained comparable proportions of all cell strains. (**Fig 5D-F and Fig S5B**). After 30 days of doxycycline treatment, non-targeting control cells dominated the tumors (69.9%), whereas triple knockdown cells were almost absent (0.06%); double knockdowns remained at ≤1.2%, and single knockdowns reached 2.2% for shAR, 5.3% for shPR, and 20.4% for shER (**Fig 5E and Table S3**).

Applying the same no-dox-normalized, NT-relative analysis to the eight-genotype competition assay (**Fig 5F and Table S3**) confirmed and extended this pattern: single-knockdown populations were reduced to 10.1% ± 0.8% (ER), 4.7% ± 1.2% (AR), and 6.0% ± 0.7% (PR) of the NT population (mean ± SEM), whereas double- and triple-knockdown populations were further depleted, to 1.9% (ER+AR), 0.7% (ER+PR), 0.1% (AR+PR), and 0.1% (ER+AR+PR) of NT. Because this normalization accounts for possible differences in baseline engraftment among the eight barcoded populations, it supports the conclusion that the additional suppression observed with combined knockdowns reflects a genuine, additive effect of targeting multiple hormone receptors simultaneously, rather than preexisting differences in the growth potential of the individual barcoded lines. Together, these results suggest a marked discrepancy in AR function between *in vitro* and *in vivo* conditions in ER-positive breast cancer and demonstrate additive growth suppression upon combined hormone receptor knockdown *in vivo*, illustrating that pooled multiplexed perturbation is a powerful strategy for functional screening while substantially reducing animal use.

## Discussion

This study presents a comprehensive and flexible lentiviral platform that overcomes several long-standing limitations in genetic perturbation technologies, particularly in the context of hormone-sensitive breast cancer (HSBC) (McGuire et al, 1977; Fannon et al, 2001; Kohler et al, 2015; Horn & Vatten, 2017). By integrating inducible RNA interference and CRISPR/Cas9-based genome editing with a range of fluorescent and luminescent reporters, we provide a scalable solution for studying gene function both *in vitro* and *in vivo*. To our knowledge, compared to existing vector systems, our platform improves multiplexing capacity, temporal control of gene expression and integration with multiparametric experimental readouts (Shaner et al, 2004; Campeau et al, 2009).

Several Gateway-based and CRISPR/Cas9 lentiviral resources are already in wide use, but each targets a different application scale. The Gateway-compatible entry/destination system underlying our overexpression module(Campeau et al, 2009) supports versatile recombination cloning for a single expression cassette, but it was not designed to combine overexpression, within one inducible platform, nor does it support native fluorescent barcoding or epitope-tags without further cloning. The Broad Institute Genetic Perturbation Platform, co-developed by Doench and colleagues, provides machine-learning-optimized sgRNA sequences (Rule Set 2/Azimuth) and genome-wide pooled lentiviral libraries for knockout (Brunello), interference (Dolcetto), and activation (Calabrese) screens (Doench et al, 2016; Sanson et al, 2018). These resources are optimized for large-scale, single-modality, constitutively expressed screening constructs assembled by restriction/Golden Gate cloning rather than Gateway recombination, and generally lack integrated fluorescent reporters or dual inducible control of Cas9 and gRNA expression. Our platform is therefore complementary to these genome-scale discovery tools: It is designed for hypothesis-driven, combinatorial, multiplexed perturbation studies with tight temporal control and validated *ex vivo* and *in vivo* applications.

A key innovation of this platform is the dual inducible architecture, which enables temporal control of gene perturbation. The use of doxycycline-regulated expression systems for both overexpression (Tet-On) and knockdown/knockout (Tet-Off for gRNA or shRNA) should ensure minimal leakage and precise activation. This is particularly valuable when studying essential genes or dynamic processes where constitutive modulation can lead to artifacts, toxicity, or loss of cell viability before the readout is achieved (Kallunki et al, 2019). The successful application of this system to USP7 and USP14 demonstrates its ability to precisely regulate Cas9 activity and achieve efficient genome editing.

Equally important is the modular design of the system, which supports a wide range of applications through the interchangeable use of fluorescent proteins, luciferases, selection markers, and functional tags. The incorporation of fourteen spectrally distinct fluorescent proteins enables the creation of unique barcoded cell populations (Lambert, 2019). These fluorescent barcodes have been shown to be uniquely resolvable in flow cytometry (with a discrimination efficiency of 95.6–98.1%) and can be used for both *in vitro* and *in vivo* multiplex analyses, reducing the need for large numbers of parallel experiments and animal cohorts.

The capacity to multiplex genetic perturbations in pooled cell populations represents a key advance. Using combinatorial knockdowns of ER, PR, and AR in MCF7 cells, we demonstrate that ER and PR knockdowns each significantly impaired proliferation and DNA synthesis, whereas knockdown of AR had negligible effects *in vitro* — consistent with its apparently limited role in MCF7 cell biology (Macedo et al, 2006). However, when these perturbations were examined *in vivo*, AR knockdown resulted in lower tumor burden. This discrepancy highlights the importance of microenvironmental cues and systemic hormonal influences in modulating gene function (Sflomos et al, 2016; Fiche et al, 2019; Constantin et al, 2025). Possible underlying mechanisms could be stromal interactions or immunomodulation, which are not present in cell cultures. These context-specific phenotypes emphasize the utility of *in vivo* validation and the potential pitfalls of relying solely on *in vitro* systems (Zhang et al, 2024).

Our barcoded competition assays *in vivo* provided further insight into receptor dependencies. By pooling all eight knockdown genotypes in individual tumors and tracking their relative abundance using flow cytometry, we demonstrated that the triple knockdown (ER+PR+AR) was strongly selected against compared to single or non-targeting controls. This competitive depletion not only confirmed the additive effects of combined receptor loss but also underscored the robustness of our multiplexing approach to assess genetic interactions in a unified biological context. Importantly, these experiments reduced the use of animals by up to eight-fold, in line with the 3Rs principle of animal research — replacement, reduction and refinement (Tannenbaum & Bennett, 2015).

A recurring concern with any pooled, barcoded competition assay is that the fluorescent tag, lentiviral integration site, or clonal origin of each population could itself influence engraftment or proliferation, independent of the intended genetic perturbation, thereby confounding interpretation of relative abundance data. To address this, we first normalized each doxycycline-treated population to its own untreated (−Dox) baseline, then expressed it relative to the equivalently normalized NT population within the same tumor (**Fig 5C, F**). Because this procedure accounts for population-specific differences in baseline growth and engraftment, the resulting ratios more directly isolate the effect of hormone receptor knockdown. The close agreement between this normalized analysis and the raw endpoint percentages (**Fig 5E**) argues against a substantial contribution of barcode- or construct-specific fitness effects and supports the conclusion that depletion of ER, PR, and AR knockdown populations, and its marked enhancement under combinatorial knockdown, reflect a genuine dependence on hormone receptor signaling rather than a technical artifact of the multiplexing strategy. More broadly, we propose this −Dox-normalized, control-relative approach as a general strategy for future pooled competition assays, in which controlling for population-specific baseline variation is essential for robust quantitative interpretation.

The utility of the system is further enhanced by biologically relevant reporters, including Fucci cell cycle probes and high sensitivity luciferases, which provide real-time readouts of cell proliferation, viability, and cell cycle status. In particular, the Fucci module enables direct observation of how specific genetic manipulations affect regulatory cell cycle transitions, leading to a deeper understanding of tumor cell dynamics and therapeutic vulnerabilities, both important aspects in cancer biology and therapy resistance (Sakaue-Sawano et al, 2008; Bajar et al, 2016).

Despite its strengths, there are also some limitations. Although our barcoding strategy works well in flow cytometry, it relies on instruments that can resolve nine or more fluorescence channels. In laboratories that do not have such instruments, multiplexing capacity may be limited. Another consideration is the scalability of this system for genome-wide applications. While our current setup is optimized for targeted perturbation studies, the design facilitates potential adaptation for ordered or pooled CRISPR and shRNA libraries (Adamson et al, 2016). In conjunction with high-content imaging or single-cell RNA-seq evaluations, this platform could be used for large-scale functional genomics screens in physiologically relevant cancer models (Maity et al, 2023; Pacesa et al, 2024).

Although this study focused on nuclear hormone receptor biology in breast cancer, the system is also transferable to other disease models. Its modularity and robust performance in both immortalized and primary cells make it well suited for studies of neurodegenerative diseases, immune regulation, stem cell differentiation and therapy resistance. In particular, the ability to monitor multiple gene perturbations in the same *in vivo* environment is a powerful strategy for identifying synthetic lethal interactions and combinatorial treatment strategies.

A related limitation concerns the semi-random, integrating nature of lentiviral delivery itself. HIV-derived lentiviral vectors preferentially integrate into transcriptionally active genes and local integration hotspots, reflecting LEDGF/p75-mediated tethering to actively transcribed chromatin (Schröder et al, 2002). This genomic context can strongly influence the behavior of doxycycline-regulated Tet-On/Tet-Off cassettes. Integration-site-dependent chromatin effects and transcriptional variegation have been reported to cause substantial clone-to-clone variability in basal (“leaky”) expression of Tet-regulated lentiviral transgenes, even among cells carrying a single vector copy(Benabdellah et al, 2016). Because our system relies on polyclonal, antibiotic- or FACS-selected populations rather than single-cell-validated clones, some degree of integration-site-dependent leakiness in the noninduced state cannot be fully excluded and may contribute to the low background editing observed over extended culture (∼4% in our hands). Transposon-based delivery systems, such as piggyBac or Sleeping Beauty, display distinct genomic integration profiles relative to lentivirus(de Jong et al, 2014) and doxycycline-inducible piggyBac constructs have, in some contexts, achieved tighter regulation with undetectable basal leakiness *in vivo*(Saridey et al, 2009), potentially reflecting the absence of viral LTR-associated regulatory elements and the lentiviral bias toward active chromatin. Incorporating chromatin insulators, safe-harbor-targeted integration, or transposon-based delivery could further reduce leaky background expression in future iterations of this platform, particularly for applications with very low tolerance for premature Cas9/gRNA activity.

These findings should be regarded as a proof-of-concept, generated in a single breast cancer cell line (MCF7) using a limited number of animals and a small panel of shRNAs whose individual potency was not exhaustively benchmarked. In particular, the finding that AR contributes to tumor growth specifically *in vivo* is based on RNA interference alone, without orthogonal rescue experiments or mechanistic follow-up, and should be interpreted as preliminary and context-specific to this model rather than as a generalizable feature of AR biology in HSBC. Broader validation across additional cell lines, PDX models, and complementary loss-of-function approaches will be needed before these observations can be extended beyond the present system.

In summary, we present a next-generation genetic perturbation platform that combines precise, inducible modulation, high-content phenotyping, and *in vivo* multiplexing within a single, modular system. By enabling the combinatorial and dynamic interrogation of genetic networks, this system enhances our capacity to model complex biological processes and may help inform future preclinical strategies for hormone-sensitive breast cancer and beyond, pending validation in additional models and cell lines.

## Materials and Methods

### Chemical reagents and antibodies

Primary antibodies used were; anti-USP7 (Bethyl, rabbit polyclonal, A300-033), USP14 (Santa Cruz, mouse monoclonal, SC-100630), anti-Cas9 (Proteintech, rabbit polyclonal, 26758-1-AP), anti-V5 (Thermo Fisher Scientific, mouse monoclonal, R960-25), anti-ERα (Santa Cruz, mouse monoclonal, sc-8002), anti-PR (Invitrogen, rabbit monoclonal, MAS-16393), anti-AR (Millipore, MP-0680, rabbit), anti-GAPDH (Abcam, rabbit polyclonal, ab9485) and anti-β-actin (Millipore, mouse monoclonal, MAB1501). For HRP-based detection, cross-adsorbed goat anti-mouse IgG-HRP (Dako, P0447) and goat anti-rabbit IgG-HRP (Dako, P0448) were used. For quantitative fluorescence imaging, IRDye 800CW goat anti-rabbit IgG (LI-COR) and IRDye 680RD goat anti-mouse IgG (LI-COR) were applied. Chemiluminescent signals were developed using Amersham ECL Prime (Cytvia).

Doxycycline hydrochloride was dissolved in ultrapure water to a stock solution of 2mg/mL and used at 1 or 0.75µg/mL. Dihydrotestosterone (DHT), promegestone (R-5020), aphidicolin and nocadazole, was kept as a stock solution in DMSO at 1µM, estradiol was diluted in EtOH to a stock solution of 1µM. All reagents were purchased from Sigma-Aldrich/Merck except for promegestone that was purchased from MedChemExpress and 5-ethynyl-2’-deoxyuridine (EdU) from Thermo Fisher Scientific.

### Cell cultures

U-2 OS were cultured in McCoys 5A + GlutaMAX Media supplemented with 1% Penicillin-Streptomycin and 10% FBS. MCF7 and HEK293T were cultured in DMEM high glucose GlutaMAX media supplemented with 1% Penicillin-Streptomycin and 10% FBS. All cells were obtained from the ATCC. All culture media and supplements were purchased from Thermo Fisher Scientific. Cell cultures were maintained at 37°C with 5% CO2 in a humidified incubator. Stably transduced inducible shRNA and gRNA cells were created by infecting target cell lines with lentiviral vectors, according to the protocol first described in (Dull et al, 1998).

### Cloning development and organization of plasmids

All the following enzymes and buffers were acquired from Thermo Fisher. For all restriction digest techniques, enzymes from the FastDigest product line were used. FastAP, T5 Ligase and Ligase Buffer were used for ligation. For PCR in preparation of sub-cloning, Phusion High-Fidelity PCR Master Mix was used, and for diagnostic PCR, DreamTaq PCR Master-Mix was used.

All plasmids, their cloning methodologies, bacterial strains used, templates and primer sequences are supplied in **Table S4**. Creation of restriction-site flanked inserts was performed either with PCR or subcloning from vectors or synthesized genes or annealing of oligonucleotides.

Oligonucleotide annealing of shRNA, gRNA and smaller tags (3XHA, Flag) was performed through 0.5°C/s cooling from 95°C. For all origins of synthesized sequences, oligonucleotides, and DNA sequences used as PCR-templates, see **Table S4**.

All plasmids were either transformed into Top10, DH5Alpha, or Stbl3 *Escherichia coli* cells. Stbl3 cells were used to prevent recombination while propagating plasmids containing LTR repeats, shRNAs, or other repetitive sequences. PureYield Plasmid MiniPrep Kit (Promega) was used for all DNA amplification steps.

Plasmid validations were performed using one or two of three methods: diagnostic restriction digest, diagnostic colony-PCR directly on bacterial colonies, and through Sanger Sequencing with Eurofins Genomics (Germany). When newly synthesized DNA was cloned, such as from oligo annealing, Sanger sequencing was always employed to verify sequence integrity.

Plasmid archiving and storage was organized through an internal identifier system. All plasmids were stored as glycerol stocks in a pre-designated location for easy retrieval and as one or more copies of miniprepped DNA for backup and use in the lab. Plasmids were never retransformed and always re-inoculated from their glycerol stock to avoid any possibility of mishandling or mutation.

### Lentivirus production and transduction

To achieve stable integration of expression constructs, lentiviral particles were generated by co-transfecting sub-confluent HEK293T cells with third-generation packaging plasmids; Gag-Pol, Rev, and VSV-G envelope (Addgene plasmids #12251, #12253, #12259) alongside either the desired expression vector or inducible shRNA/gRNA plasmids. Transfections were carried out using the calcium phosphate method. Viral supernatants were harvested at 48- and 72-hours following transfection. For transduction, target cells were exposed to a 1:1 mix of viral supernatant and fresh complete medium, supplemented with 8 μg/mL polybrene. After 48 hours, cells were re-plated at low density and selected with either FACS (for mTagBFP2 transduction) or with antibiotics; blasticidin (8 μg/mL) for 7 days, puromycin (2 μg/mL) for 3–5 days, hygromycin B (200 μg/mL) for 7–10 days, or neomycin/G418 (500 μg/mL) for 10–12 days. Selection media were refreshed every three days during the selection period.

For the lentiviral titration experiment lentivirus carrying a doxycycline-inducible Cas9-V5 construct was produced in HEK293T cells as described above and concentrated ∼18-fold. Physical titer was determined by p24 ELISA (VPK-107, Cell Biolabs) and converted to an estimated functional titer using an empirical correction factor for large-insert packaging efficiency of 1:1000. Reported MOIs (10–70) are therefore nominal physical MOIs, estimated functional MOIs are 10-100-fold lower. U2OS cells were transduced at each nominal MOI and selected with blasticidin. Stable lines were treated with doxycycline and Cas9 expression quantified by RT-qPCR. Cas9 induction after 48h doxycycline was similar across all selected MOIs, indicating that antibiotic selection enriches for transduced cells regardless of starting MOI, making constructs as large as inducible Cas9 expression vectors, largely independent of viral dose within the range of titers tested (**Fig S6**). All shRNA and gRNA expression plasmids used in this study were of comparable size, and no consistent differences in packaging or transduction efficiency were observed between vectors. Titer was more strongly influenced by plasmid purity and preparation than by insert identity. Because all stable lines were generated as polyclonal populations, selected by antibiotic resistance or FACS rather than isolated as single-cell clones, we assumed that lentiviral integration occurred comparably across these lines and relative to the non-targeting control. We therefore did not perform karyotyping or vector copy number analysis on individual lines. Knockdown efficiency, assessed by Western blot, was comparable among single, double, and triple knockdown lines **(Fig 4E–G)**, indicating that combinatorial transduction with multiple shRNAs did not measurably compromise silencing efficiency for each target.

### Relative quantitative PCR

For validation of Cas9 expression, cells were retrieved from plates with Trizol and extracted with Direct-Zol RNA MiniPrep kit (Zymo Research). cDNA was synthesized with iScript Advanced cDNA synthesis kit (BioRad). qPCR was performed and analyzed with CFX 96 (BioRad). For qPCR primers utilized, see **Table S4**.

### Estimating editing efficiency of USP7 edit in inducible Cas9 pools

The frequency of edited alleles was estimated by amplicon sequencing using Oxford Nanopore Technologies (ONT). Briefly, genomic (g)DNA of edited cells was extracted using a MagMAX DNA Multi-Sample Ultra 2.0 Kit and a KingFisher Duo Prime instrument (both ThermoFisher Scientific). The isolated gDNA samples were then subjected to PCR using primers to amplify the targeted region in USP7 (5’- ACACTCTTTCCCTACACGACGCTCTTCCGATCTTCCTCAGCCTGCCAAATAGT-3’ and 5’-GCACGTATCTTGCTGCAAACA-3’). The forward primer has an overhang which was used in a second PCR for indexing. Sequencing libraries of the PCR products were then prepared using the ONT Native Barcoding Kit 24 V14, following the manufacturer’s instructions. Sequencing was performed using a MinION Mk1D device and accompanying flow cell (ONT) with real-time base calling using MinKNOW software. Obtained reads were aligned to a reference sequence using minimap2 and filtered using SAMtools to retain only primary alignments with a mapping quality score (MAPQ) ≥ 20, excluding unmapped, secondary, and supplementary alignments. Editing events (insertions/deletions) surrounding the guide target site (10 bp upstream and downstream of the guide binding site) were detected and used to calculate the frequency of edited alleles. Sequencing results are presented in **Table S1**.

### *In vitro* luciferase assay

Expression of NanoLuc and AkaLuc was confirmed by *in vitro* luciferase assay. Cells were seeded in white 96-well plates (Greiner) at 4,000 cells/well in complete medium. The following day, the medium was replaced with phenol red-free DMEM supplemented with 200 µM luciferase substrate Furimazine for NanoLuc or Akalumine-HCl for AkaLuc and luminescence was immediately measured using a 470 ± 40 nm filter for NanoLuc or an open filter setting for AkaLuc on a CLARIOstar microplate reader (BMG Labtech).

### *In vivo* Bioluminescence Imaging

Bioluminescence imaging was performed using the Xenogen IVIS Imaging System 200 (Caliper Life Sciences). Mice were injected intraperitoneally with D-luciferin (150 mg/kg in PBS; Biosynth) and anesthetized with isoflurane 10 minutes post-injection. Images were acquired with automatic exposure settings using Living Image software (Caliper Life Sciences). For *ex vivo* imaging of metastases, 450 mg/kg luciferin was administered 7 minutes prior to euthanasia. Signal quantification was performed by measuring total photon flux (photons/second) in manually drawn regions of interest (ROIs) around mammary glands or organs.

### Animal Studies

NOD.Cg-*Prkdc^scid^ Il2rg^tm1Wjl^*/SzJ (NSG) mice were purchased from Charles River Laboratories. All animal procedures were approved by the Service de la Consommation et des Affaires Vétérinaires of Canton de Vaud, Switzerland (authorization VD 3795), and carried out in accordance with Swiss animal welfare guidelines. Mice were housed under specific pathogen-free (SPF) conditions with a 12-hour light/dark cycle, constant temperature (22 ± 2 °C), and humidity (55 ± 10%). Animals had ad libitum access to food and acidified water (pH 2.5–3.0) in polysulfone bottles. Mice were kept in IVC Green Line cages (Type II long; Tecniplast) with aspen bedding (Tapvei), cardboard houses, and wooden tunnels for enrichment. Mice were fed with either standard chow diet (Provimi-Kliba, cat. #3242) or doxycycline-supplemented chow (0.625 g/kg; Safe, E8404P01R 00069), replaced weekly.

### Intraductal Cell Injection

Intraductal injection of cells was performed as previously described (Sflomos et al, 2016). Female NSG mice (8–12 weeks) were anesthetized via intraperitoneal injection of ketamine (75 mg/kg) and xylazine (10 mg/kg). Once anesthetized, ophthalmic ointment was applied to prevent corneal dehydration. Fur around the 3rd and 4th mammary glands was shaved, and the area was disinfected. Under a stereomicroscope, nipples were trimmed to expose the ductal orifice, which was visualized using 2% trypan blue and gently wiped. An equal mixture of fluorescently barcoded cell populations, totaling 200,000 single cells in 10 µL of sterile PBS, was injected into each mammary gland using a 33-gauge blunt-end Hamilton syringe (HAMI80508). Mice received 0.5 mL of 0.9% saline intraperitoneally post-injection for rehydration and were monitored on a heating pad until recovery. Paracetamol (200–300 mg/kg) was administered in drinking water (500 mg in 250 mL) starting one day before and continuing for three days after the procedure to minimize discomfort.

### Tissue Dissociation and Flow Cytometry

Isolated mammary gland tissues mechanically cut into small fragments (0.3–0.5 mm) using two sterile scalpels. The chopped tissue was transferred into a gentleMACS™ C Tube (Miltenyi Biotec) containing 5 mL of dissociation medium and processed on the gentleMACS™ Octo Dissociator with heating block using the program 37°C_h_TDK_2. After completion of the program, the tissue suspension was gently resuspended by pipetting up and down with a 5 mL pipette. 10 mL of PBS were added, and the sample was centrifuged at 300g for 5 min. The supernatant was discarded, and the pellet was resuspended in 5 mL of trypsin solution, followed by incubation at 37 °C for 5 min. Subsequently, 5 mL of PBS supplemented with FBS (PBS/FBS) and 100 µL of DNase I (10 mg/mL) were added, and the mixture was incubated at 37 °C for 5 min before centrifugation at 300g for 5 min. The resulting pellet was resuspended in 5 mL of red blood cell (RBC) lysis buffer and incubated for 5 min at room temperature. After RBC lysis, 10 mL of PBS/FBS were added, and the suspension was centrifuged again at 300 × g for 5 min.

For flow cytometry analysis cells were resuspended in FACS buffer (PBS containing 2% FBS and 1 mM EDTA), filtered through a 40 µm strainer, and kept on ice. Fluorescent reporters (mTagBFP2, eGFP, iRFP670, sfCherry3C) were analyzed using single-color controls for spectral compensation. Gating strategies were defined using single color samples, negative control samples from uninjected mammary glands or glands injected with wild-type (WT) cells. Flow cytometry was performed on a BD LSRFortessa, and cell sorting was carried out using a BD FACSAria II or MoFlo Astrios cell sorter. Flow cytometry data were analyzed using FlowJo software (BD Biosciences). Gating strategy described in **Fig S2**.

For multiplex fluorescence experiments, cells were cultured in complete growth medium, harvested by trypsinization, washed twice with 10% FBS/PBS, and resuspended in 5% FBS/PBS. Cells were filtered through a 35 μm mesh and kept on ice prior to sorting. Nine-color cell sorting was performed using a BD FACSymphony S8 (BD Biosciences). Compensation was applied using single-color controls. Sorted cells were collected into 10% FBS/PBS and assessed for post-sort purity. FlowJo software (BD Biosciences) was used for analysis. Gating strategy described in **Fig S1**.

### Viability Assays

#### Resazurin assay

Cells were seeded in 96-well black-walled, clear-bottom plates at 2 × 10^4^ cells/well in complete growth medium supplemented with doxycycline (0.75 µg/mL). After 24 hours, medium was replaced with phenol red–free, charcoal-stripped DMEM containing doxycycline. On day 4, medium was refreshed with the same stripped medium supplemented with doxycycline and 10 nM each of dihydrotestosterone (DHT), estradiol, and promegestone. On day 7 (three days of hormone treatment), cell viability was assessed using a resazurin reduction assay. A working solution of resazurin (0.02 mg/mL) was prepared by diluting a 4 mg/mL stock 1:200 in complete medium and added directly to each well (100 µL/well) without media removal. Plates were incubated at 37 °C for 4–5 hours, and fluorescence was measured using a ClarioSTAR plate reader (Ex: 560 nm, Em: 590 nm). Viability was normalized to control-treated wells.

#### Cell confluency measurements

Cells were plated at a density of 2,500 cells per well in 96-well plates containing regular growth medium. doxycycline was added on day 0 to a final concentration of 1 μg/mL and refreshed on day 3. Parallel wells without doxycycline treatment served as controls. Plates were transferred to the Incucyte SX5 live-cell imaging system (Sartorious), and images were automatically acquired at defined time intervals over a period of 7 to 10 days. Cell confluence was quantified over time and used as a measure of proliferation.

#### EdU staining

Cells were seeded in 96-well black-walled, clear-bottom plates at 1.6-2.5 × 10^4^ cells/well in complete growth medium supplemented with doxycycline (0.75 µg/mL). On day 1, the medium was replaced with phenol red–free, charcoal-stripped FBS supplemented DMEM containing doxycycline. On day 2, compounds were added at 10 nM final concentration via serial dilution (1:100 from stock, then 1:10 into media). On day 3, cells were pulsed with 10 µM EdU (5-ethynyl-2′-deoxyuridine; 10 mM stock in DMSO) for 30 minutes at 37 °C. Cells were then fixed in 4% paraformaldehyde for 15 minutes at room temperature, permeabilized with PBS containing 0.1% Triton X-100 for 15 minutes. EdU detection was performed using a homemade click chemistry cocktail freshly prepared in PBS. The reaction mix contained 4 mM CuSO₄, 2 µM azide-conjugated fluorophore (ATTO-488), and 10 mM ascorbic acid, added in that order to initiate the reaction. After removing PBS, 100 µL of reaction mix was added per well, and plates were incubated for 30 minutes at room temperature, protected from light. Following staining, cells were washed three times with PBS and counterstained with DAPI (1µM for 10 min at RT). Images were acquired using CellcyteX imaging system (Echo). Image analysis was performed using CellProfiler with segmentation of nuclei based on DAPI and cell cycle data was analysed in R. Data were normalized to control-treated wells.

### Western Blot

For USP7 and USP14 depletion confirmation, cells were harvested by washing once in PBS and cell pellets were lysed in RIPA buffer containing Mini EDTA-free protease inhibitor cocktail (Roche) on ice for 20 min with intermittent pipetting. Cell lysates were centrifuged at 12 000 x *g* and 4°C for 20 min. The supernatant containing the total cellular protein was quantified by BCA assay (Thermo Scientific) and denatured at 95°C for 5 mins using Laemmli sample buffer (Bio-Rad). Proteins were separated on Any kD™ Mini-PROTEAN® TGX™ precast gels (Bio-Rad) and Precision Plus Protein™ Dual Color standards (Bio-Rad) were used as ladders for molecular weight comparisons. Separated proteins were transferred using Trans-Blot® Turbo™ transfer system (Bio-Rad) on to nitrocellulose membranes (Bio-Rad) and blocked with Intercept (TBS) Blocking Buffer (LI-COR) for one hour at room temperature. Primary antibodies; rabbit anti-USP7 (1:7,500), mouse anti-USP14 (1:500), rabbit anti-Cas9 (1:3000), mouse anti-β-actin (1:3,000) and rabbit-GAPDH (1:2,500) were prepared in Intercept (TBS) Blocking Buffer and incubated overnight with the membranes at 4°C. Membranes were washed three times with TBS + 0.05% Tween-20 (TBST) and incubated with LI-COR IRDye® secondary antibodies diluted in prepared Intercept (TBS) Blocking Buffer or ROTI®Stock TBST (Carl Roth) (1:10,000) for 1h at room temperature in the dark. Finally, membranes were washed three times with TBST and imaged using a LI-COR Odyssey® Fc imaging system. All Western blots were analyzed and quantified using Image Studio software.

For experiments validating knockdown of hormone receptors, cells were washed in ice-cold PBS and lysed in 1% SDS buffer or RIPA buffer with protease inhibitors (Merck). Lysates were incubated on ice for 20 minutes and sonicated using a Bioruptor Plus (Diagenode) for 15 cycles (20 s ON/20 s OFF, high power), followed by a centrifugation at 12000 g for 20 min. Protein concentration was determined using the BCA assay (Thermo Scientific) and measured with a SpectraMax iD3 plate reader (Molecular Devices). Samples were denatured at 95 °C for 5 minutes and stored at −20 °C. 40-50 µg of protein was resolved on 12% SDS-PAGE gels and transferred to nitrocellulose membranes using the Trans-Blot system (Bio-Rad). Membranes were blocked in 5% milk/TBS-T for 1 hour and incubated overnight at 4 °C with primary antibodies diluted in blocking buffer. Primary antibodies used were mouse anti-ER (1:400), rabbit anti-PR (1:400), rabbit anti-AR (1:1,000) and mouse anti-β-actin (1:3,000). After washing, membranes were incubated with HRP-conjugated secondary antibodies (at 1:5,000–1:10,000) for 1 hour at room temperature. Signals were detected using ECL substrate (Amersham, Cat. No. RPN2232) and imaged on a ChemiDoc system (Bio-Rad). For re-probing, membranes were stripped with 0.1 M glycine buffer (pH ∼2.5) for 20 minutes and re-blocked prior to re-incubation with antibodies.

Antibody specificity in the experiments used was supported by the expected reduction or loss of the corresponding protein signal following target-specific knockout or knockdown.

### Immunofluorescence and quantification of Fucci reporters

Fluorescence images in Fig 3C-D and Fig 4C were acquired using an EVOS M5000 Imaging System (Invitrogen) equipped with 10× phase-contrast objective (EVOS FL PH, NA 0.30). Live cell images represented in Fig 3C-D were acquired after incubation of indicated compounds for 24 hours. Fucci reporter signals were quantified per nucleus in CellProfiler using the upper-quartile intensities of mClover and mKO2 fluorescence. Cell-cycle-like states were assigned using integrated Hoechst intensity as a DNA-content proxy, with thresholds defined from DMSO controls and applied unchanged to all treatment groups.

### Statistical Analysis

Statistical analyses and graph generation were performed using GraphPad Prism version 10. Specific details regarding statistical methods, sample sizes, and significance values are included in each figure legend, for the *in vivo* statistics, the doxycycline treated animal/samples was normalized towards the corresponding average of the doxycycline untreated animal/sample for cell numbers. ANOVA with Fishers LSD post-hoc towards NT were run as a statistical test. Exact p-values are indicated in each graph.

## Data Availability

All data supporting the findings of this study are provided within the article and its supplementary materials. Additional original data are available from the corresponding author upon reasonable request.

## Acknowledgments

The authors gratefully acknowledge the MedH Core Flow Cytometry Facility at Karolinska Institutet, supported by KI/SLL, for expert assistance with flow cytometry, and the SciLifeLab CRISPR Functional Genomics Unit at Karolinska Institutet for expert support with Oxford Nanopore Technologies (ONT) sequencing.

In addition, we would like to thank Danny Labes and Mariela Castelblanco from the AGORA Flow Cytometry Facility for their expert assistance with cell sorting and flow cytometry analyses, and Soohyun Moon for technical assistance with Western blot experiments.

## Author contributions

Conceptualization (MA), Methodology (SA, DM, JB, MJP, NCKV, AP, SE, KS, HQ, MA), Validation (JB, SA, MJP, DM), Formal analysis (SA, NCKV MJP, DM), Investigation (SA, MJP, DM, JB), Resources (NCKV, MA, CB), Writing – original draft (SA, JB, DM), Writing – review & editing: (SA, MA, NCKV, CB), Visualization (JB, SA, MJP, DM, MA), Supervision (NCKV, MA, CB), Project administration (MA), Funding acquisition (NCKV, MA, CB).

## Supplementary Figures

**Supplementary Figure 1.**
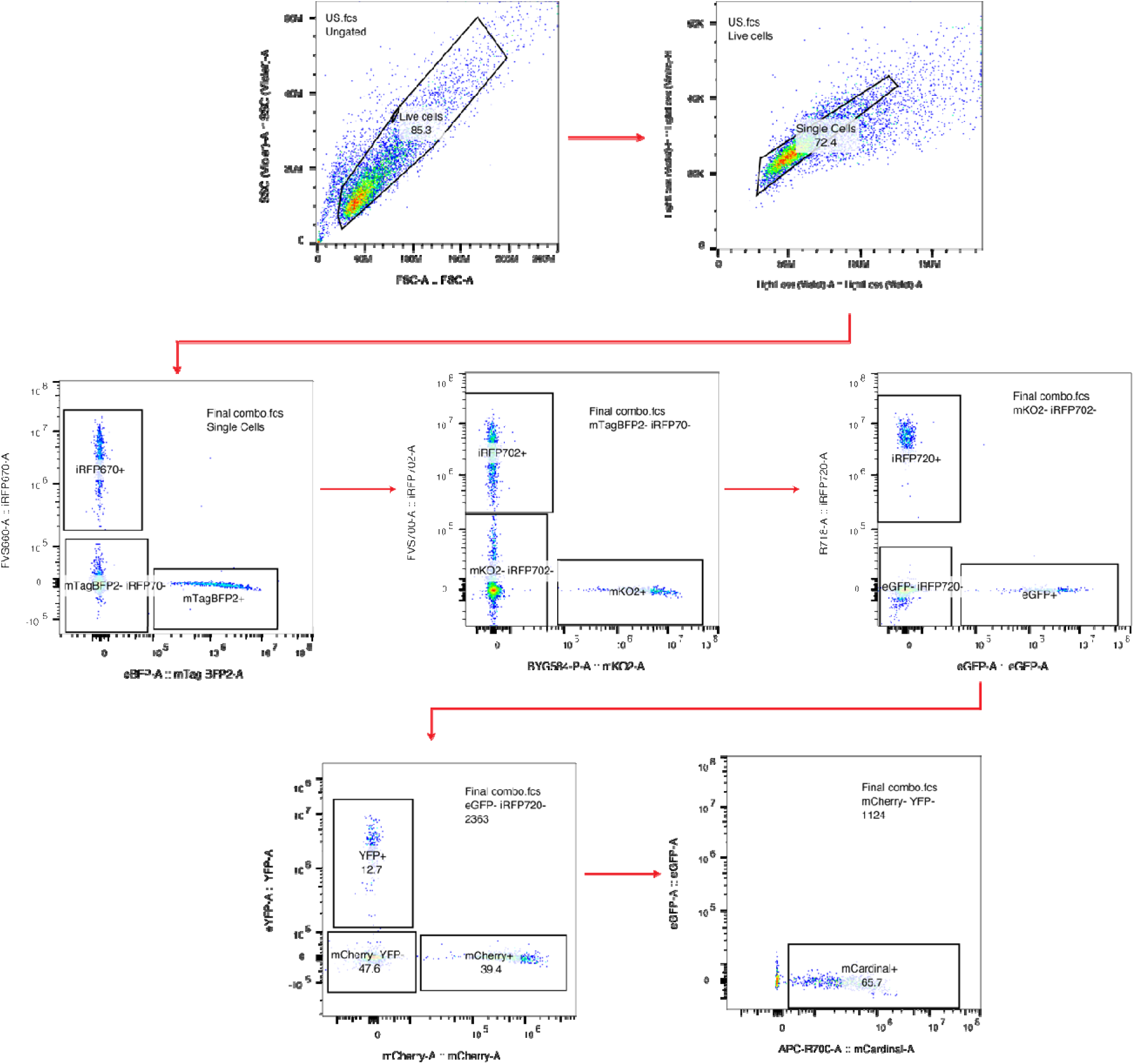
Gating strategy of multicolor FACS experiment of single-color barcoded cells. Live cells were gated for exclusion of debris and doublets to identify single cells. Cells within this population were then gated to separate each fluorophore expressing cell in the following order; mTagBFP2 vs iRFP670, mKO2 vs iRFP702, eGFP vs iRFP720, mCherry vs YFP and mCardinal.

**Supplementary Figure 2.**
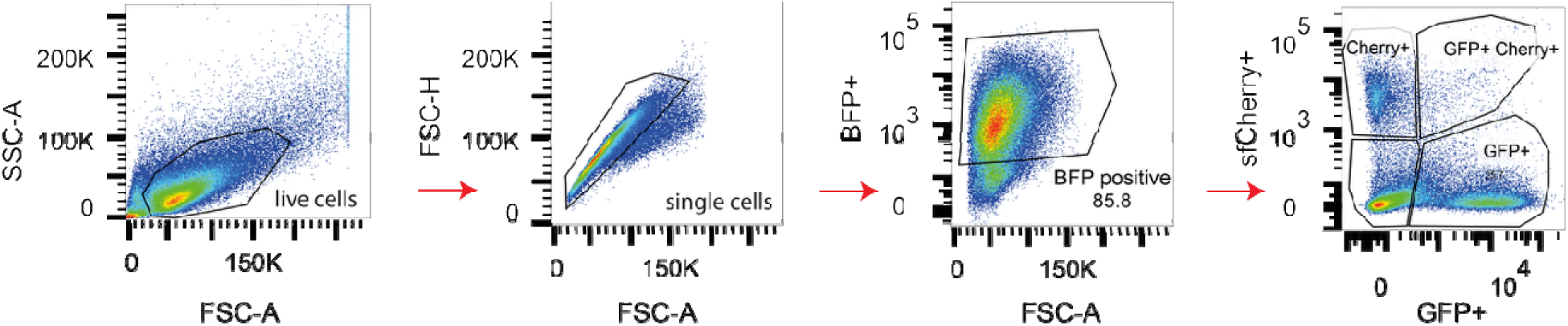
Flow cytometry gating strategy used to identify and quantify fluorescently barcoded MCF7 cell populations. Representative gating strategy illustrating the sequential selection of MCF7 cell populations isolated from MIND model. Cells were first gated for live cells (FSC-A vs. SSC-A) and singlets (FSC-H vs. FSC-A). Within the singlet population, mTagBFP2□ cells were selected, and subsequent gating was applied to identify additional fluorescent markers. Subpopulations expressing GFP and/or sfCherry3C within the mTagBFP2□fraction (mTagBFP2□eGFP□, mTagBFP2□sfCherry3C□, and mTagBFP2□eGFP□sfCherry3C□) are shown here as an example. The same gating strategy was applied to identify and quantify iRFP670-expressing cells and to determine the proportion of single-positive mTagBFP2□cells.

**Supplementary Figure 3.**
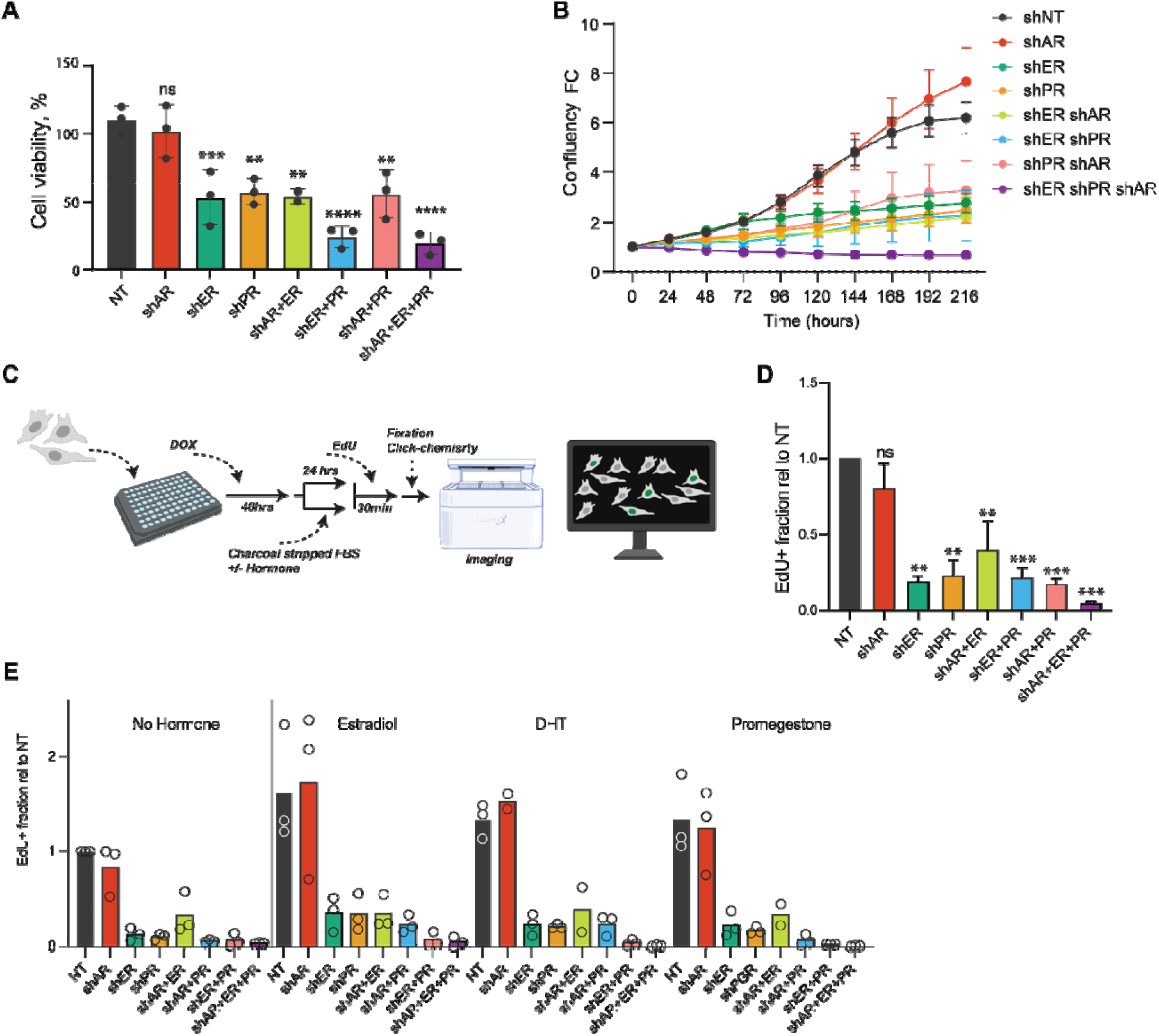
Effects of single, double, and triple hormone receptor knockdowns on MCF7 proliferation and DNA synthesis. **(A)** Resazurin-based metabolic viability assay after 8 days of doxycycline induction in cells with knockdowns targeting ER, PR, and AR individually or in combinations. Data normalized to untreated controls and represent mean ± SD of three biological replicates. Statistical analysis was performed using one-way ANOVA with Dunnett’s multiple comparisons test. **(B)** Proliferation curves monitored over 10 days by live-cell imaging (IncuCyte), shown as fold change relative to time zero. Data represent mean ± SEM from two independent experiments. **(C)** Schematic workflow for EdU incorporation assay including cell seeding, doxycycline treatment, hormone supplementation, EdU labeling, click chemistry, and imaging using CellCyte imager. **(D)** Quantification of EdU-positive fraction relative to non-targeting control under standard growth conditions. Data represent mean ± SD from at least three experiments. **(E)** EdU incorporation results in charcoal-stripped FBS media supplemented with DOX, vehicle, estradiol, dihydrotestosterone (DHT), or promegestone. Data normalized to NT and represent mean ± SD, from two independent experiments.

**Supplementary Figure 4.**
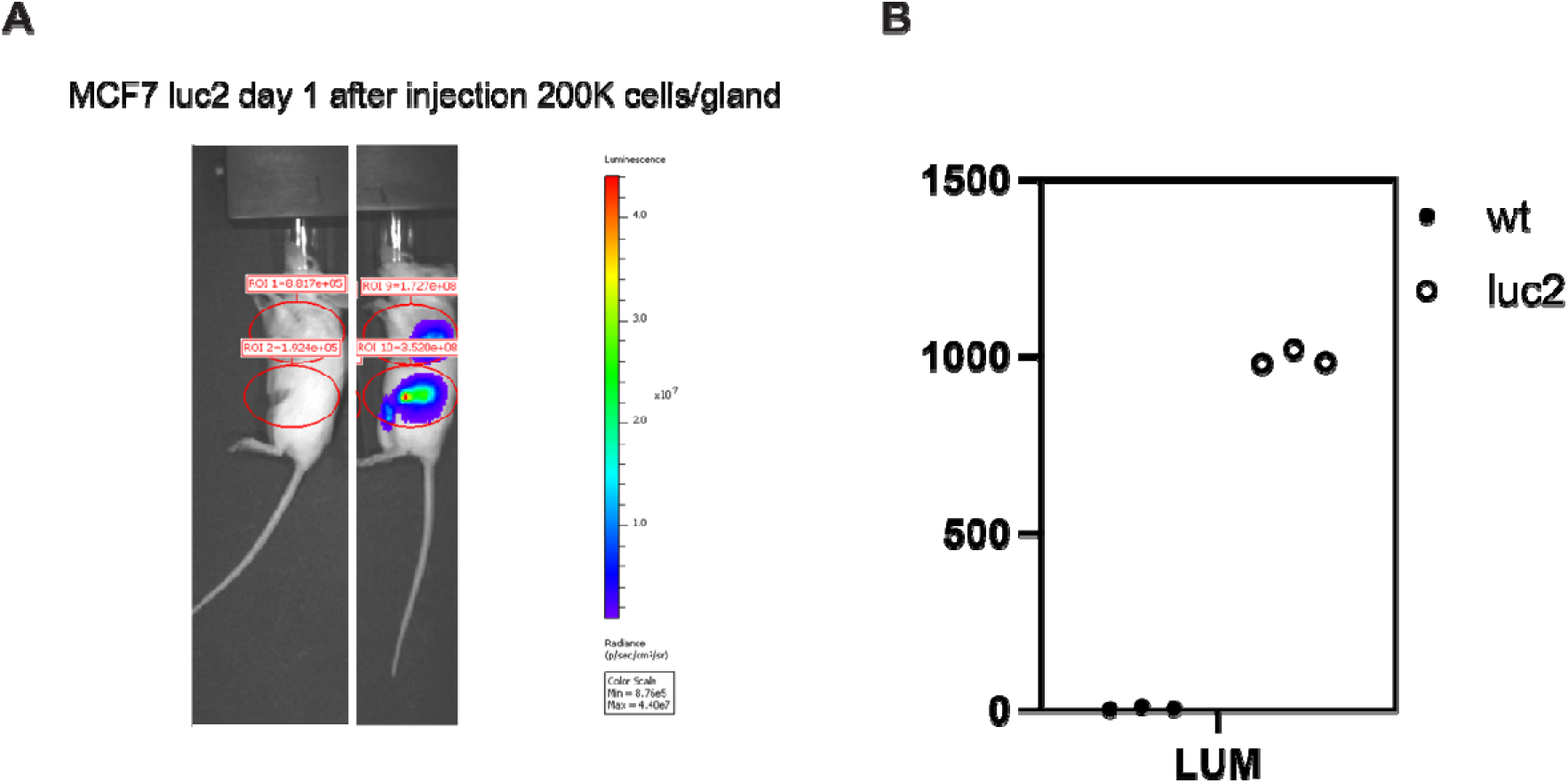
*In vivo* and *in vitro* bioluminescence imaging of MCF7-RFP-Luc2 cells. (A) Representative IVIS image showing *in vivo* bioluminescence signal in the third and fourth mammary glands of an NSG mouse injected intraductally with 200,000 MCF7-Luc2 cells, one day after injection. Regions of interest (ROIs) corresponding to each injected mammary gland are indicated. The color scale represents photon flux (photons/sec/cm²/sr). (B) *In vitro* measurement of bioluminescence intensity in MCF7-Luc2 cells compared with wild-type (WT) MCF7 cells, confirms specific luciferase activity in MCF7-RFP-Luc2 cells relative to WT controls.

**Supplementary Figure 5.**
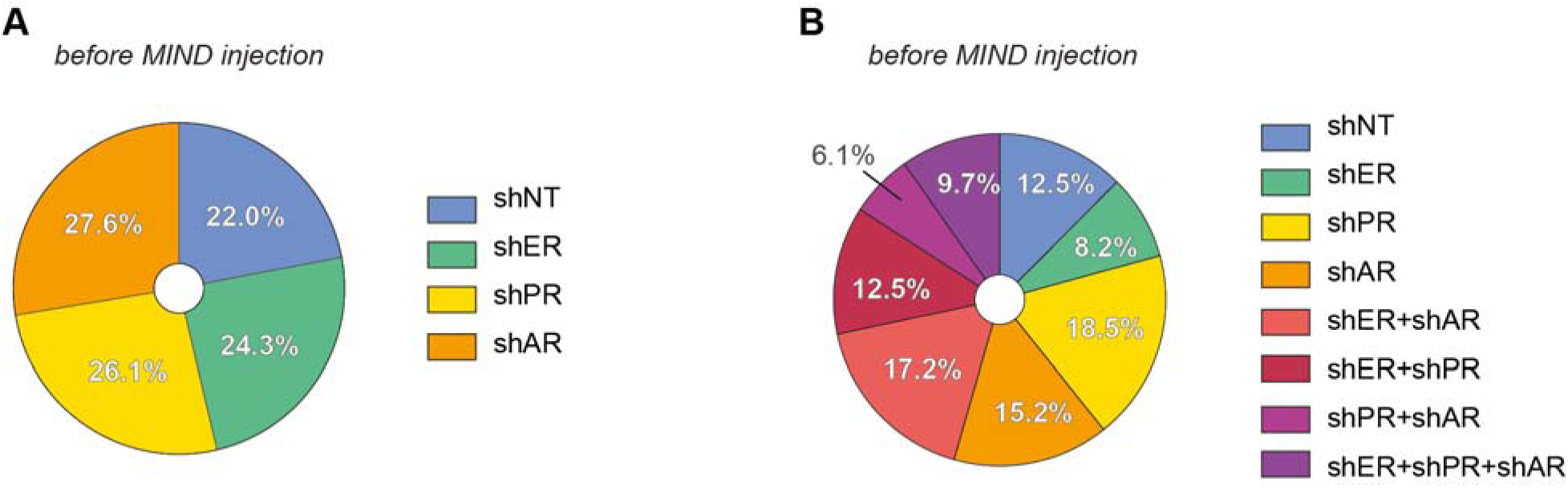
Distribution of fluorescently labeled populations in the pooled cell mixture before injection into mice. Fluorescently labeled populations in the pooled cell mixture before injection were measured by flow cytometry. Each color represents a distinct knockdown condition, where (**A**) shows the single knockdowns of four labeled MCF7 cell lines and (**B**) shows the full panel of eight different cell lines. The distributions are shown as percentages of the total injected population.

**Supplementary Figure 6.**
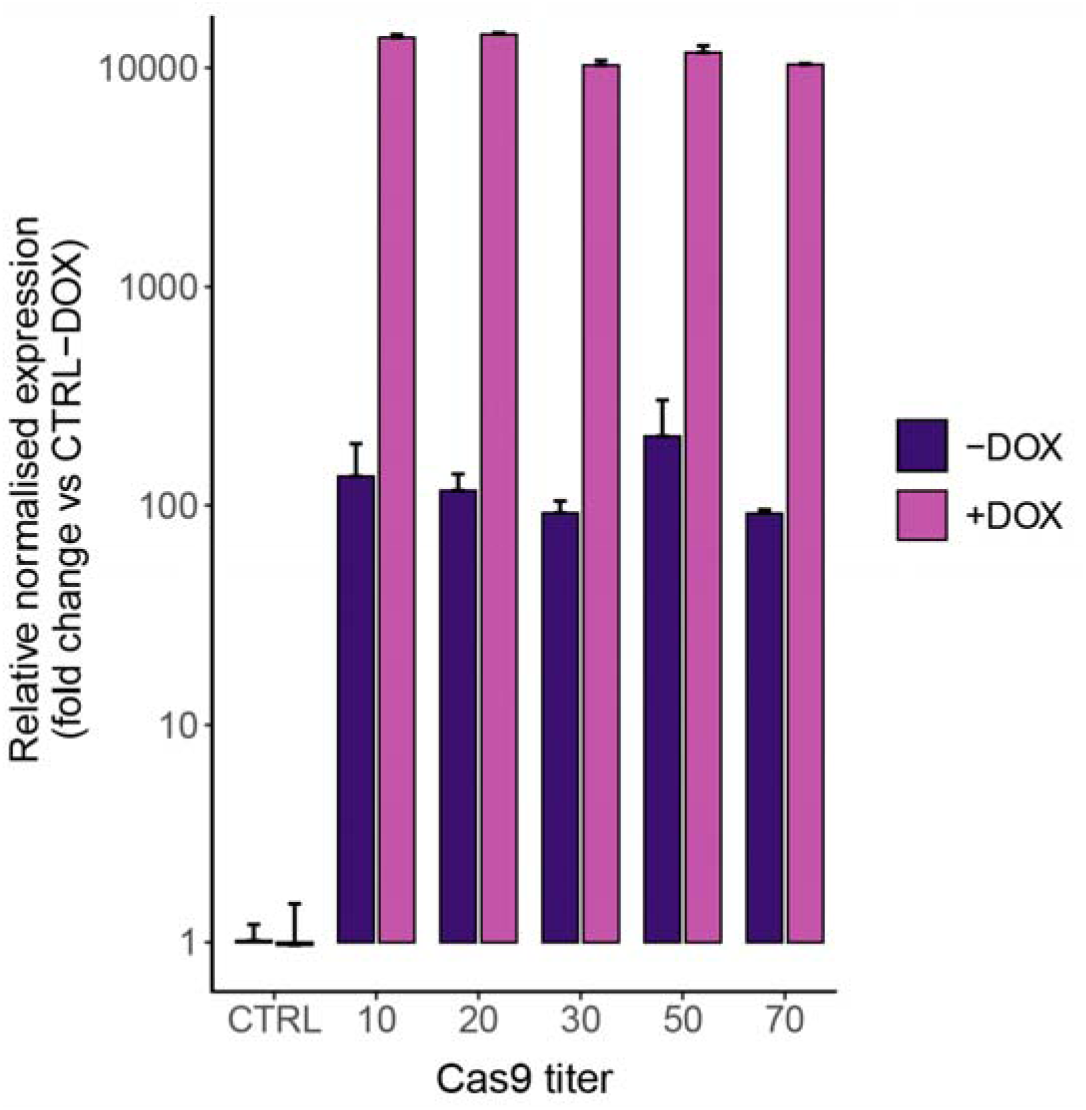
Virus titer did not affect inducible Cas9 expression after selection of transduced cells. After blast selection, the number of cells obtained correlated with the virus titer, but after expansion and Cas9 induction we did not detect any significant differences among the Cas9-expressing cell lines by qPCR. Basal Cas9 expression measured by qPCR is 100-fold higher than primer-dimer amplification observed in untransduced control cells. After Cas9 induction with doxycycline, Cas9 expression is 100-fold higher than basal levels.

